# Extension of PERMANOVA to Testing the Mediation Effect of the Microbiome

**DOI:** 10.1101/2022.04.26.489586

**Authors:** Ye Yue, Yi-Juan Hu

## Abstract

**Background:** Recently, we have seen a growing volume of evidence linking the microbiome and human diseases or clinical outcomes, as well as evidence linking the microbiome and environmental exposures. Now comes the time to assess whether the microbiome mediated the effects of the exposures on the outcomes, which will enable researchers to develop interventions to modulate the outcomes by modifying the microbiome composition. Use of distance matrices is a popular approach to analyzing complex microbiome data that are high-dimensional, sparse, and compositional. However, the existing distance-based methods for mediation analysis of microbiome data, MedTest and MODIMA, only work well in limited scenarios.

**Results:** PERMANOVA is currently the most commonly used distance-based method for testing microbiome associations. Using the idea of inverse regression, here we extend PERMANOVA to testing microbiome mediation effects by including both the exposure and the outcome as covariates and basing the test on the product of their *F*-statistics. This extension of PERMANOVA, which we call PERMANOVA-med, naturally inherits all the flexible features of PERMANOVA, e.g., allowing adjustment of confounders, accommodating continuous, binary, and multivariate exposure and outcome variables including survival outcomes, and providing an omnibus test that combines the results from analyzing multiple distance matrices. Our extensive simulations indicated that PERMANOVA-med always controlled the type I error and had compelling power over MedTest and MODIMA. Frequently, MedTest had diminished power and MODIMA had inflated type I error. Using real data on melanoma immunotherapy response, we demonstrated the wide applicability of PERMANOVA-med through 16 different mediation analyses, only 6 of which could be performed by MedTest and 4 by MODIMA.

**Availability and Implementation:** PERMANOVA-med has been added to the existing function “permanovaFL” in our R package LDM, which is available on GitHub at https://github.com/yijuanhu/LDM.

## Background

Microbiome research has proliferated in the last decade due to booming interests in the scientific community, increasing power of high-throughput sequencing, and rapid advancement of data analytics. So far, most microbiome studies have been focused on bivariate associations between the microbiome and the covariates of interest. We have seen a rapidly growing volume of evidence linking the microbiome and human diseases such as obesity [1] or clinical outcomes such as response to immunotherapy [2]. Similarly, we have seen the relationship between the microbiome and environmental exposures such as diet [3]. Meanwhile, many of these environmental exposures have well-established effects on the clinical outcomes. We believe now comes the time to assess whether the microbiome play a mediating role between the exposures and the outcomes, as depicted in Figure 1(a). Identifying such a mediating role of the microbiome enables researchers to develop interventions to modulate the outcomes by modifying the microbiome composition. Accordingly, there is an urgent need for statistical methods are designed specifically for mediation analysis of microbiome data.

**Figure 1:**
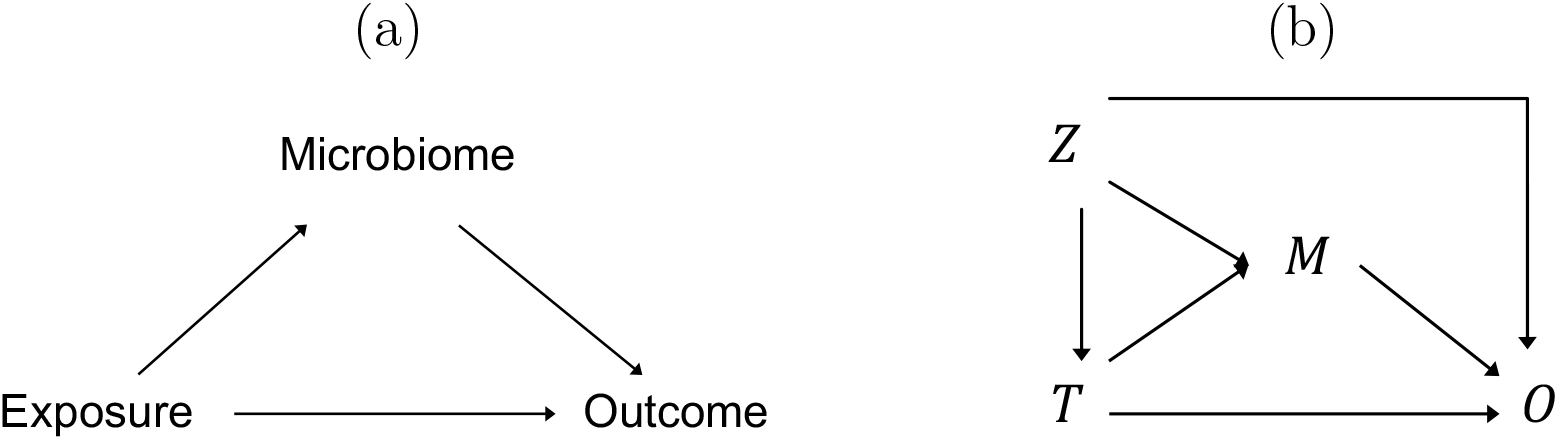
(a) Some effect of the exposure on the outcome is mediated through the microbiome. (b) *T* denotes the exposure, *M* the microbiome (mediator), *O* the outcome, and *Z* the confounding covariates.

There are a number of special challenges in mediation analysis of microbiome data. Microbiome read count data from 16S amplicon or metagenomic sequencing are typically summarized in a taxa count table, which have unique and complex features. They are high-dimensional (with typically many more taxa than samples), sparse (having 50–90% zero counts), compositional (measuring relative abundances that sum to one), and highly overdispersed. In addition, microbiome studies can be conducted in various observational or clinical settings, and tend to have diverse attributes. The exposure and outcome variables may be continuous, binary, or even multivariate (comprising multiple components such as multiple indicators for a categorical variable). In particular, many clinical outcomes are in the form of times to event (survival times) with possibly censored values. There generally exist confounders (e.g., sex and antibiotic use), as a microbial community is easily modifiable. Finally, the sample sizes are usually small (e.g., 50–100) and the study designs can be complex (e.g., clustered samples [4] or matched sets [5]).

To circumvent the complexities of microbiome count data, a popular approach is to first summarize the taxon-level data into a distance (dissimilarity) matrix that measures the pairwise dissimilarity in the microbiome profiles, and then base the analysis of microbiome data on the distance matrix [6–10]. This approach provides results at the community level, which is usually the first step in an analytical pipeline. Numerous distance measures, with different properties, have been proposed to detect diverse patterns in microbiome data; the most commonly used ones include Jaccard [11], Bray-Curtis [12], and weighted or unweighted UniFrac [13, 14]. It is well acknowledged that the optimal choice of a distance measure depends on the underlying variation pattern in a particular dataset, which is unknown a priori. Therefore, it is a common practice to construct an omnibus test that combines the results from analyzing different distance matrices.

Two existing methods, MedTest [15] and MODIMA [16], adopted such a distance-based approach to mediation analysis of microbiome data. Specifically, MedTest uses the principal components (PCs) of a given distance matrix as multiple mediators and tests their joint mediation effects. However, the assumption that the exposure-microbiome association and the microbiome-outcome association coincide at the same set of PCs may be overly optimistic. Also, the PCs may not capture mediation effects at rare taxa. Moreover, MedTest does not accommodate multivariate exposures and outcomes in its current form. MODIMA calculates distance matrices from the exposure, the microbiome, and the outcome, separately, and employs the distance correlation [17, 18] for characterizing the exposure-microbiome association and the partial distance correlation [19] for the microbiome-outcome association conditional on the exposure. The distance matrices for the exposures and outcomes naturally accommodate multivariate variables. However, MODIMA does not allow adjustment of confounders and does not provide an omnibus test. Finally, neither MedTest nor MODIMA can handle censored survival times.

PERMANOVA [7] is currently the most commonly used distance-based method in analysis of microbiome data. Although it was originally developed for testing microbiome associations, we find that we can extend PERMANOVA to testing microbiome mediation effects by using the idea of inverse regression and including both the exposure and the outcome as covariates whose *F*-statistics capture the exposure-microbiome association and the microbiome-outcome association conditional on the exposure, respectively. This extension of PERMANOVA would naturally inherit all the features of PERMANOVA, some of which have been a focus of recent development, including adjustment of confounders [4], test of multivariate covariates, test of censored survival times [20], and an omnibus test of multiple distance matrices [21]. Thus, the extension of PERMANOVA would be very appealing to researchers who routinely use PERMANOVA.

In this article, we present PERMANOVA-med, the extension of PERMANOVA to testing the community-level mediation effect of the microbiome. We base PERMANOVA-med on our implementation of PERMANOVA through the function “permanovaFL” in our R package LDM [4], which differs from the “adonis2” implementation in the R package vegan in the permutation scheme and outperformed adonis2 in many situations [4, 5, 22]. In the methods section, we first motivate the use of inverse regression and then show how to extend PERMANOVA to PERMANOVA-med. In this process, we provide an overview of PERMANOVA, as well as overviews of MedTest and MODIMA to facilitate comparison with PERMANOVA-med. In the results section, we present extensive simulation studies in which we numerically compared PERMANOVA-med to MedTest and MODIMA. We also demonstrate the wide applicability of PERMANOVA-med through 16 different mediation analyses of the real data on melanoma immunotherapy response. We conclude with a discussion section.

## Methods

### Motivation towards inverse regression

The relationships among the exposure (*T*), mediator (*M*), outcome (*O*), and confounders (*Z*) are depicted in Figure 1(b). Assuming a continuous outcome and a continuous mediator and further assuming no exposure-mediator interaction and no unmeasured confounding, the classical mediation model [23] specifies a linear model for the mediator and a linear model for the outcome:

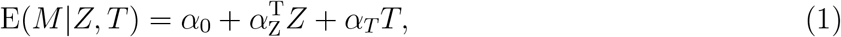

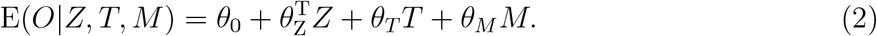

Note that *α*_*T*_ characterizes the effect of *T* on *M* given *Z*, and *θ*_*M*_ characterizes the effect of *M* on *O* given *Z* and *T*. Then it can be shown that the mediation effect is given by *α*_*T*_ *θ*_*M*_ [24]. However, it is unclear how to use the microbiome composition data, which are represented by a distance matrix here, as a mediator. Also, the *forward* outcome model (2) is not easily generalizable to an outcome variable that is discrete, multivariate, or censored survival time. These limitations motivated us to adopt the *inverse* regression model that exchanges the positions of the outcome and the mediator in model (2). Inverse regression is a commonly used approach to testing associations [25–27]. It has a key advantage of accommodating different types of outcome variables including multivariate variables. In what follows, we show that, by proper orthogonalization of the non-microbiome variables, the inverse regression model we consider “merges” both models (1) and (2) into one regression model, which fits nicely into the framework of PERMANOVA that takes the distance matrix as the response variable.

To be specific, we first sequentially orthogonalize variables *Z, T*, and *O*, and denote the residual of *T* after orthogonalizing against *Z* by *T*_*r*_ and denote the residual of *O* after orthogonalizing against (*Z, T*) by *O*_*r*_. Then, we consider the inverse regression model

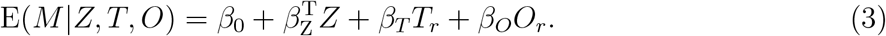

For now, we view *M* as a univariate continuous variable, just as in (1) and (2). Model (3) implies that 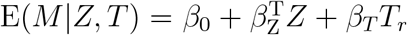, which is exactly model (1) after replacing *T* by *T*_*r*_. Thus, we easily obtain that *β*_*T*_ = *α*_*T*_. Although it is well known that *β*_*O*_ ≠ *θ*_*M*_, we see that *β*_*O*_ = 0 and *θ*_*M*_ = 0 coincide as they both capture the microbiome-outcome association given (*Z, T*). As a result, testing *β*_*T*_ *β*_*O*_ = 0 is equivalent to testing *α*_*T*_ *θ*_*M*_ = 0, i.e., whether there exists a mediation effect through *M*. We find that model (3) fits nicely into the PERMANOVA framework, in which we view *M* as a distance matrix and the linear regression as a partition of *M* into additive components corresponding to the orthogonal factors (*Z, T*_*r*_, *O*_*r*_).

### Overview of PERMANOVA

PERMANOVA is based on a linear model of covariates that partition a given distance matrix along each covariate. In particular, when the Euclidean distance measure is used on the relative abundance data, it is the total variance of relative abundance data across all taxa that is partitioned into variance explained by each covariate. Following our implementation in permanovaFL [4], we denote the design matrix of all covariates by *X* and group the columns of *X* into *K* submodels, i.e., *X* = (*X*_1_, *X*_2_, …, *X*_*K*_). Each submodel includes components that will be tested jointly, such as a single covariate, multiple covariates, or multiple indicators for a categorical covariate. The submodels are first processed into sequentially orthogonal, unit vectors by the Gram-Schmidt process, so that the partition of the distance matrix is unambiguous. This requires that the covariates in *X* follow a scientifically meaningful order; for example, the confounders should enter first. Let *D* denote the *n* × *n* distance matrix calculated among *n* samples, which is often Gower-centered [28] to become Δ = −0.5 (*I* − *n*^−1^11^′^) *D*^2^ (*I* − *n*^−1^11^′^), where *D*^2^ is the element-wise squared *D, I* is the identity matrix, and 1 is a vector of *n* ones. The “residual” distance matrix after projecting off all submodels except the *k*th one takes the form 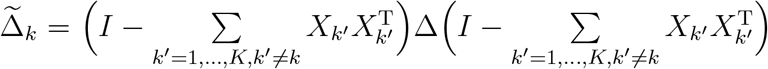by noting that 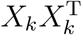 is the hat matrix for the *k*th submodel. Then, PERMANOVA tests the effect of the *k*th submodel by using the *F*-statistic

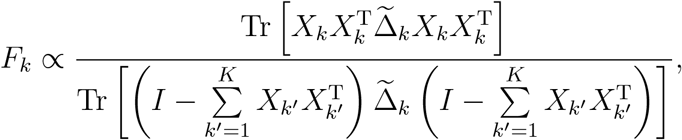

where Tr(·) is the trace operation. PERMANOVA assesses the significance of the *F*-statistic via permutation, particularly the Freedman-Lane permutation scheme [29] as implemented in permanovaFL. The Supplementary Materials of [4] showed that the Freedman-Lane scheme is equivalent to forming the following statistic for the *b*th permutation replicate:

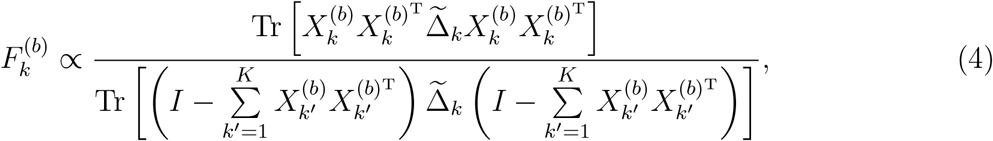

where 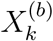 is a row-permuted version of *X*_*k*_ and thus the columns of 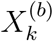 remain orthogonal. Note that the residual distance matrices 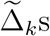 do not need to be recalculated for each replicate. In contrast, the permutation scheme implemented in adonis2 replaces all 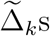 in *F*_*k*_ and 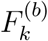 by the raw distance matrix Δ.

PERMANOVA is very versatile. It can handle censored survival times. As proposed in [20], the survival times and censoring statuses are first fit by a Cox model (including non-microbiome risk predictors as covariates) to be converted into the Martingale or deviance residuals, which are then used as a generic continuous covariate in PERMANOVA. Because PERMANOVA bases its inference on permutation, it is robust to small sample sizes. The permutation replicates can also be readily used to construct an omnibus test of multiple distance matrices, which uses the minimum of the *p*-values obtained from analyzing each distance matrix as the final test statistic and uses the corresponding minima from the permutation replicates to simulate the null distribution. In addition, the permutation can be conducted in ways that preserve the correlation found in the original data, so PERMANOVA can accommodate certain structures of samples such as clustered samples [4] and matched sets [5]. All the features that PERMANOVA supports were summarized in Figure 2.

**Figure 2:**
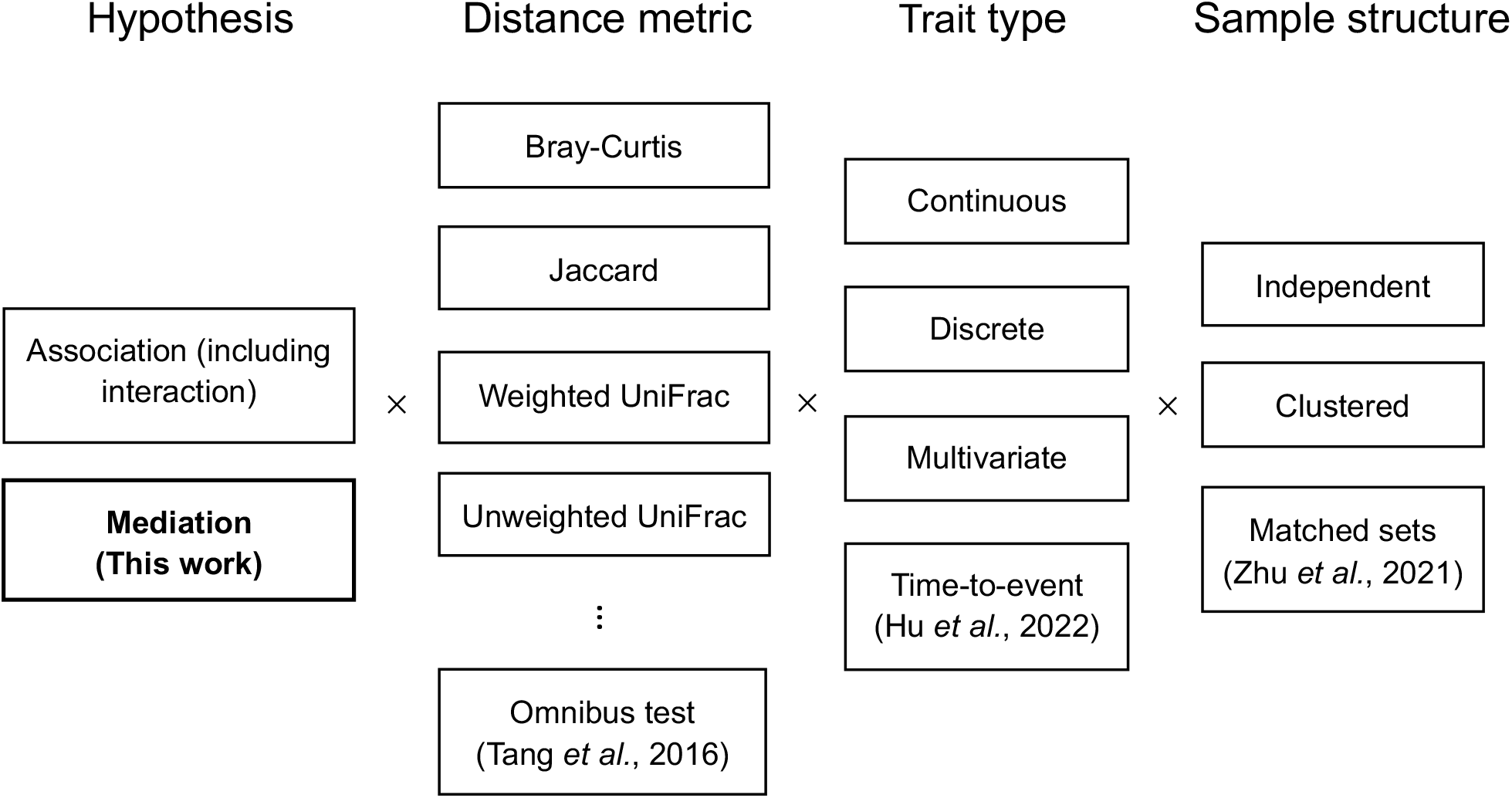
Analyses supported by permanovaFL. Analysis types without a citation were introduced in the original LDM paper [4]. “Clustered” refers to analyses of clustered data where traits of interest vary by cluster or vary both by and within clusters (some analyses may require special structure or additional assumptions). “Matched sets” is a special type of clustered data in which all traits of interest vary within sets.

### Extension of PERMANOVA to mediation analysis

Under model (3), we set submodels *X*_1_ = *Z, X*_2_ = *T*_*r*_, and *X*_3_ = *O*_*r*_ and denote the PERMANOVA *F*-statistics for testing microbiome associations with *T*_*r*_ and *O*_*r*_ by *F*_*T*_ and *F*_*O*_, respectively. Then, we propose to test the existence of a mediation effect by the microbiome, i.e., *H*_0_ : *β*_*T*_ *β*_*O*_ = 0, using the test statistic

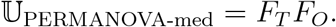

To claim a mediation effect by the microbiome, both the exposure-microbiome and microbiome-outcome associations (given the exposure) are required to be significant. Thus, the null hypothesis of no mediation is a composite null that consists of no exposure-microbiome association, no microbiome-outcome association, or neither. Accordingly, we construct the statistic for the *b*th permutation replicate,

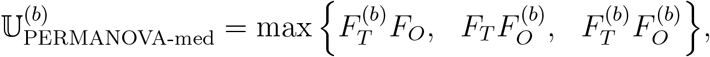

where the three product terms correspond to the statistics under the three types of null hypotheses. Then, the *p*-value is obtained as the proportion of 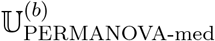 that are equal to or larger than the observed statistic U_PERMANOVA-med_. Note that all the *F*-statistics needed for calculating the *p*-value are directly available from PERMANOVA. As a result, our mediation analysis implemented in the PERMANOVA framework naturally inherits all the features in PERMANOVA.

### Overview of MedTest and MODIMA

MedTest considers microbiome “features” to be the eigenvectors of the Gower-centered distance matrix Δ, denoted by *u*_1_, *u*_2_, …, *u*_*L*_, that are associated with the *L* positive eigenvalues, denoted by *λ*_1_, *λ*_2_, …, *λ*_*L*_. MedTest assumes that these microbiome features are the units through which the microbiome exert the mediation effect. Thus, it adopts a test statistic that is a sum of feature-specific mediation effects, each weighted by *λ*_*l*_ (the percentage of variance explained by that feature):

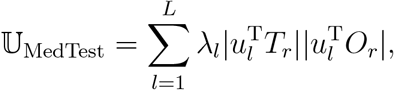

where |*·*| is the absolute value function. Note that 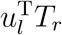 and 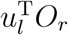 are the sample Pearson correlation coefficients that measure the associations between the *l*th feature and the exposure and the outcome, respectively; the sample Pearson correlation coefficient does not easily accommodate multivariate exposure or outcome variables. Similar to PERMANOVA-med, MedTest calculates the maximum of the statistics corresponding to the three types of null hypotheses for the *b*th permutation replicate:

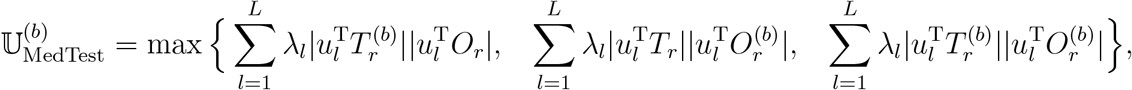

where 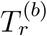 and 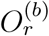 are permuted vectors of *T*_*r*_ and *O*_*r*_, respectively. Finally, the *p*-value is obtained as the proportion of 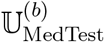 that are equal to or larger than the observed statistic U_MedTest_. The power of MedTest may critically depend on whether the exposure-microbiome association and the microbiome-outcome association coincide at the same set of PCs. Further, when the true mediators in the community are rare taxa, the PCs may not effectively capture the variation at these mediators.

In addition to the distance matrix *D* from the microbiome profiles, MODIMA also requires the *n* × *n* distance matrices (usually the Euclidean distance) being calculated from the exposure data and the outcome data, separately, which we denote by *D*_*T*_ and *D*_*O*_. These distance matrices naturally accommodate multivariate variables. Then, MODIMA uses the distance correlation [18], dCor(*D*_*T*_, *D*), for measuring the exposure-microbiome association, which parallels the Pearson correlation with the major difference being that the centered product moment transformation is applied to the distance matrices rather than data vectors. MODIMA uses the partial distance correlation [19], pdCor(*D*_*O*_, *D*|*D*_*T*_), for measuring the microbiome-outcome association conditional on the exposure, which parallels the Pearson partial correlation. MODIMA adopts the test statistic

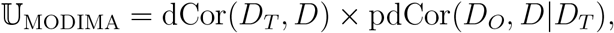

and the statistic for the *b*th permutation replicate,

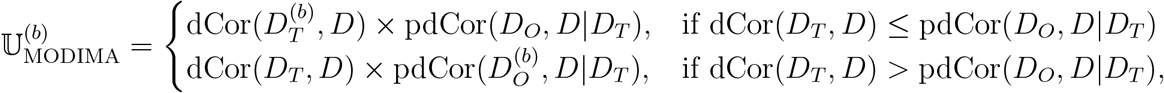

where 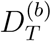 and 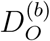 are obtained by permuting both rows and columns of the *D*_*T*_ and *D*_*O*_ matrices, respectively. Although this way of constructing the null statistic appears different from those in PERMANOVA-med and MedTest, they seem asymptotically equivalent. Finally, the *p*-value is calculated as the proportion of 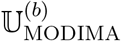 that are equal to or larger than the observed statistic U_MODIMA_. Note that, in this process, the confounding covariate *Z* cannot be adjusted. Further, the MODIMA paper pointed out a lack of correspondence between conditional independence and zero partial distance correlation, e.g., a non-zero partial correlation in scenarios with conditionally independent variables. It implies that MODIMA may generate false positive findings under the null hypothesis of no mediation, especially when there is a strong direct effect of the exposure on the outcome (*θ*_*T*_ in model (2)).

## Results

### Simulation studies

Our simulations were based on data on 856 taxa of the upper-respiratory-tract (URT) microbiome [30], and the mediator model (1) and the forward outcome model (2) as generative models. We considered both binary and continuous exposure variables, continuous outcome variables, and 100 or 200 sample size (*n*); note that both MedTest and MODIMA papers considered continuous exposures only. In what follows, we number the taxa by decreasing relative abundance so that taxon 1 is the most abundant. We considered three mediation mechanisms, in which we assumed the mediating taxa were the top five most abundant taxa (taxa 1–5), 100 relatively rare taxa (taxa 51–150), and a mixture of abundant and relatively rare taxa (taxa 4, 5, 51, and 52), which are referred to as M-common, M-rare, and M-mixed, respectively. We further assumed that the mediating taxa played the role through their relative abundances in M-common and M-mixed and through their presence-absence (0/1) statuses in M-rare.

Specifically, for a binary exposure *T*_*i*_, we assigned half of the samples *T*_*i*_ = 1 and the other half *T*_*i*_ = 0. For a continuous exposure *T*_*i*_, we sampled *T*_*i*_ from the Beta(2, 2) distribution. We initially set the baseline relative abundances of all taxa for all samples to the population means that were estimated from the real data, which we denote by 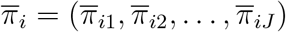. To induce the effects of the exposure on the mediating taxa, we decreased 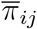 by the percentage *β*_TM_*T*_*i*_ (∈ [0, 1]) for taxa 3–5 in M-common and taxa 5 and 51 in M-mixed, and then redistributed the decreased amount evenly over the remaining mediating taxa, i.e., taxa 1–2 in M-common and taxa 4 and 52 in M-mixed. In M-rare, we set 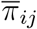 for the mediating taxa to 0 with the probability *β*_TM_*T*_*i*_ independently, and increased 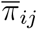 of the most abundant taxon by the total mass that had been set to 0 (which did not affect the presence-absence statuses of the most abundant taxon as it was always present). This way of modifying 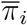 did not change the relative abundances of non-associated taxa (except for the most abundant taxon in M-rare) and the modified 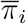 still satisfied the compositional constraint (unit sum). Note that *β*_TM_ characterizes the exposuremicrobiome (T-M) association and *β*_TM_ = 0 corresponds to no T-M association. Next, we drew the sample-specific composition *π*_*i*_ = (*π*_*i*1_, *π*_*i*2_, …, *π*_*iJ*_) from the Dirichlet distribution 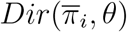 where the overdispersion parameter *θ* was set to 0.02 (as estimated from the real data). Then, we generated the read count data using the Multinomial distribution with mean *π*_*i*_ and library size (sequencing depth) sampled from *N* (10000; (10000*/*3)^2^) and truncated at 2000. Finally, we scaled each read count by the library size to obtain the observed relative abundance, denoted by *M*_*ij*_ for taxon *j* in sample *i*.

In M-common and M-mixed, we generated the continuous outcome *O*_*i*_ from the following model that allows different directions for the effects of different taxa on the outcome:

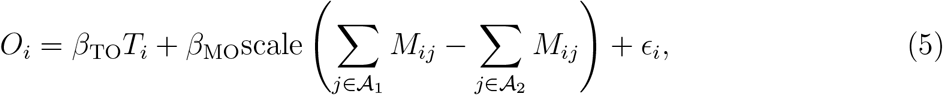

where 𝒜_1_ and 𝒜_2_ are the “increasing” and “decreasing” subsets of mediating taxa as determined above and *ε*_*i*_ ∼ *N* (0, 0.5^2^). In M-rare, we let 𝒜_1_ and 𝒜_2_ to include taxa 51–100 and taxa 101–150, respectively, and replaced *M*_*ij*_ in (5) by *I*(*M*_*ij*_ ≠ 0). We also considered a modification of the microbiome-outcome (M-O) association by restricting 𝒜_1_ and 𝒜_2_ to a subset of originally selected taxa, i.e., taxa 4 and 5 in M-common, taxa 51 and 52 in M-mixed, and taxa 101–150 in M-rare.

We simulated a binary confounder *Z*_*i*_ in settings with a binary exposure. Note that a confounder is associated with the exposure, the microbiome, and the outcome simultaneously (Figure 1(b)). First, we generated *Z*_*i*_ = 1 with probability 0.7 among samples with *T*_*i*_ = 1 and with probability 0.3 among those with *T*_*i*_ = 0. Then, we used the same operation as used for simulating the T-M association, except that we replaced *β*_TM_*T*_*i*_ by *γ*_ZM_*Z*_*i*_ with *γ*_ZM_ = 0.6, to further modify 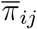 based on *Z*_*i*_ for the mediating taxa that had been modified based on *T*_*i*_. Finally, we added the term *γ*_ZO_*Z*_*i*_ with *γ*_ZO_ = 0.7 to model (5).

We applied PERMANOVA-med and compared it to MedTest and MODIMA, for testing the mediation effect of the microbiome in the simulated data. In M-common and M-mixed, all tests were based on the Bray-Curtis distance. In M-rare, all tests were based on the Jaccard distance. The type I error and power of all tests were assessed at the nominal level 0.05 based on 10000 and 1000 replicates of data, respectively.

### Simulation results

We first present results for the simulated data without a confounder. The power of the PERMANOVA-med, MedTest, and MODIMA with varying values of *β*_MO_, *β*_TM_, *β*_TO_, and sample size *n* are displayed in Figures 3, 4, and 5 for M-common, M-mixed, and M-rare, respectively. The numerical values of the type I error rates (when *β*_MO_ = 0) shown in these figures are also listed in Table 1.

**Table 1:** Type I error (at the level 0.05) in analysis of simulated data without a confounder

| Scenario | Exposure | $\beta_{TM}$ | $\beta_{TO}$ | $n$ | PERMANOVA-med | MedTest | MODIMA |
| --- | --- | --- | --- | --- | --- | --- | --- |
| M-common | Binary | 0.2 | 0.1 | 100 | 0.012 | 0.021 | 0.017 |
|  |  | 0.4 | 0.1 | 100 | 0.044 | 0.049 | 0.046 |
|  |  | 0.4 | 0.8 | 100 | 0.044 | 0.049 | 0.086 |
|  |  | 0.4 | 0.8 | 200 | 0.046 | 0.052 | 0.126 |
|  | Continuous | 0.4 | 0.1 | 100 | 0.009 | 0.016 | 0.013 |
|  |  | 0.6 | 0.1 | 100 | 0.026 | 0.032 | 0.025 |
|  |  | 0.6 | 0.8 | 100 | 0.026 | 0.032 | 0.040 |
|  |  | 0.6 | 0.8 | 200 | 0.048 | 0.045 | 0.072 |
| M-mixed | Binary | 0.4 | 0.1 | 100 | 0.014 | 0.019 | 0.017 |
|  |  | 0.6 | 0.1 | 100 | 0.039 | 0.043 | 0.040 |
|  |  | 0.6 | 0.8 | 100 | 0.039 | 0.043 | 0.047 |
|  |  | 0.6 | 0.8 | 200 | 0.048 | 0.049 | 0.068 |
|  | Continuous | 0.6 | 0.1 | 100 | 0.004 | 0.010 | 0.007 |
|  |  | 0.8 | 0.1 | 100 | 0.011 | 0.016 | 0.013 |
|  |  | 0.8 | 0.8 | 100 | 0.011 | 0.016 | 0.016 |
|  |  | 0.8 | 0.8 | 200 | 0.027 | 0.033 | 0.038 |
| M-rare | Binary | 0.2 | 0.1 | 100 | 0.039 | 0.041 | 0.042 |
|  |  | 0.4 | 0.1 | 100 | 0.050 | 0.028 | 0.041 |
|  |  | 0.4 | 0.8 | 100 | 0.050 | 0.028 | 0.088 |
|  |  | 0.4 | 0.8 | 200 | 0.052 | 0.023 | 0.125 |
|  | Continuous | 0.6 | 0.1 | 100 | 0.045 | 0.046 | 0.042 |
|  |  | 0.8 | 0.1 | 100 | 0.044 | 0.034 | 0.039 |
|  |  | 0.8 | 0.8 | 100 | 0.044 | 0.034 | 0.082 |
|  |  | 0.8 | 0.8 | 200 | 0.049 | 0.026 | 0.125 |

Note: The type I error results were generated at $\beta_{MO} = 0$ (i.e., no M-O association), and thus the same for datasets using different sets of taxa for generating the M-O association.

In M-common with a binary exposure, when the same abundant taxa (taxa 1–5) were used to generate both the T-M and M-O associations (Figure 3(a)), MedTest was slightly more powerful than PERMANOVA-med, possibly because the top PCs used by MedTest effectively captured both the T-M and M-O associations. When a subset of taxa (taxa 4 and 5) were used for generating the M-O association (Figure 3(b)), the power of MedTest declined much more quickly than the power of PERMANOVA-med, as the PCs that captured the T-M association (e.g., PC1) may not coincide with the PCs that captured the M-O association (e.g., PC2). MODIMA seemed to be very powerful in some cases (e.g., Figure 3(a)), but its performance was sensitive to the value of *β*_TO_. In particular, MODIMA generated inflated type I error when *β*_TO_ was enlarged to 0.8 and especially when *n* was also increased from 100 to 200.

**Figure 3:**
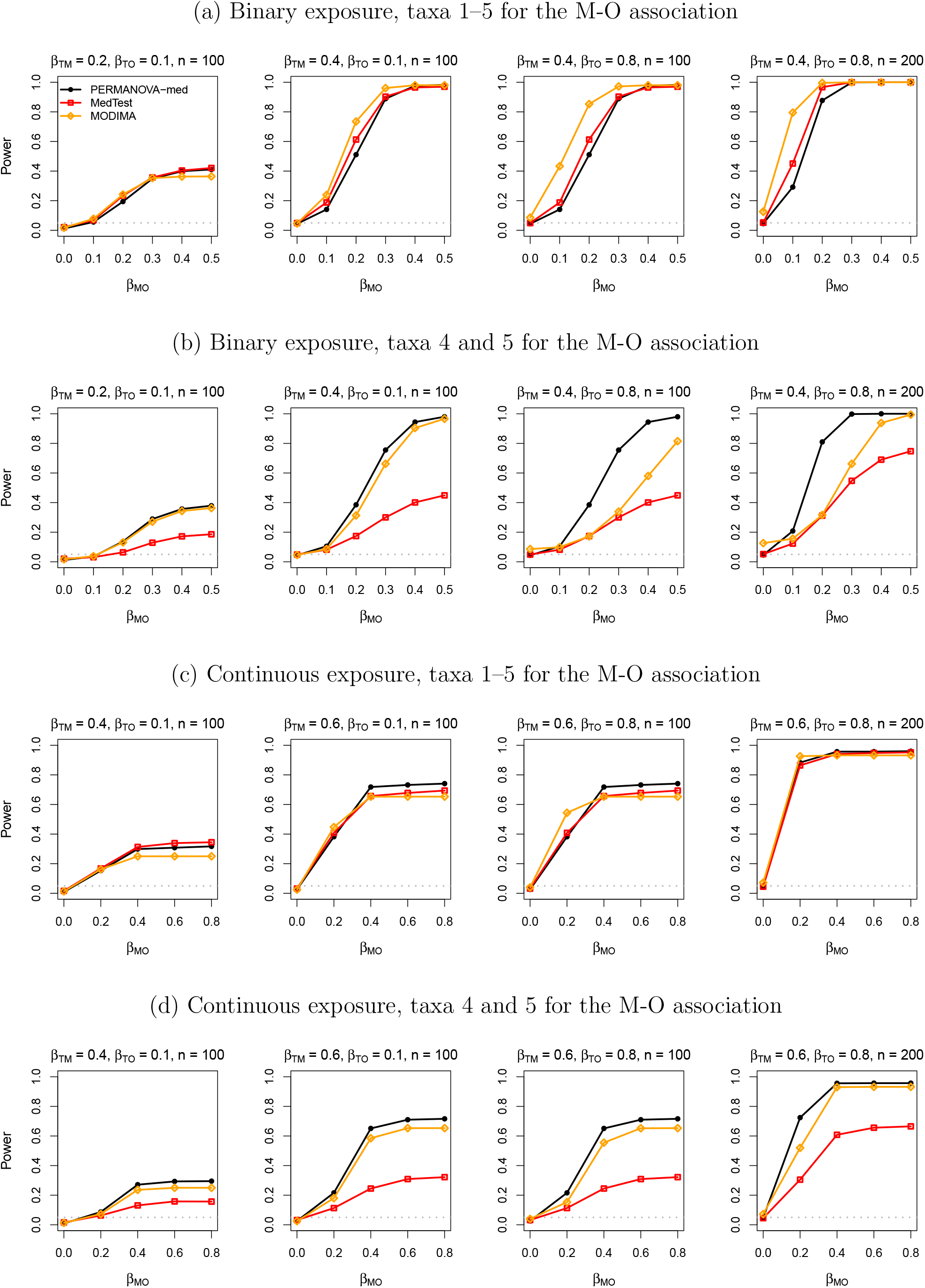
Simulation results in analysis of simulated data under M-common.

In M-common with a continuous exposure, which tended to result in more complex variation patterns in the data than a binary exposure, MedTest (and MODIMA) lost the advantage in power to PERMANOVA-med, even when taxa 1–5 were used for both the T-M and M-O associations (Figure 3(c)). Again, MedTest lost further, considerable power to PERMANOVA-med when taxa 4 and 5 were used for the M-O association (Figure 3(d)) and MODIMA yielded inflated type I error when *β*_TO_ and *n* were both large.

As expected, PERMANOVA-med always had significantly higher power than MedTest in M-mixed (Figure 4), and the power difference was more pronounced in M-rare (Figure 5), since PCs became less efficient in capturing variations in less abundant taxa. In M-rare, MODIMA was uniformly less powerful than PERMANOVA-med, even its type I error was clearly inflated.

**Figure 4:**
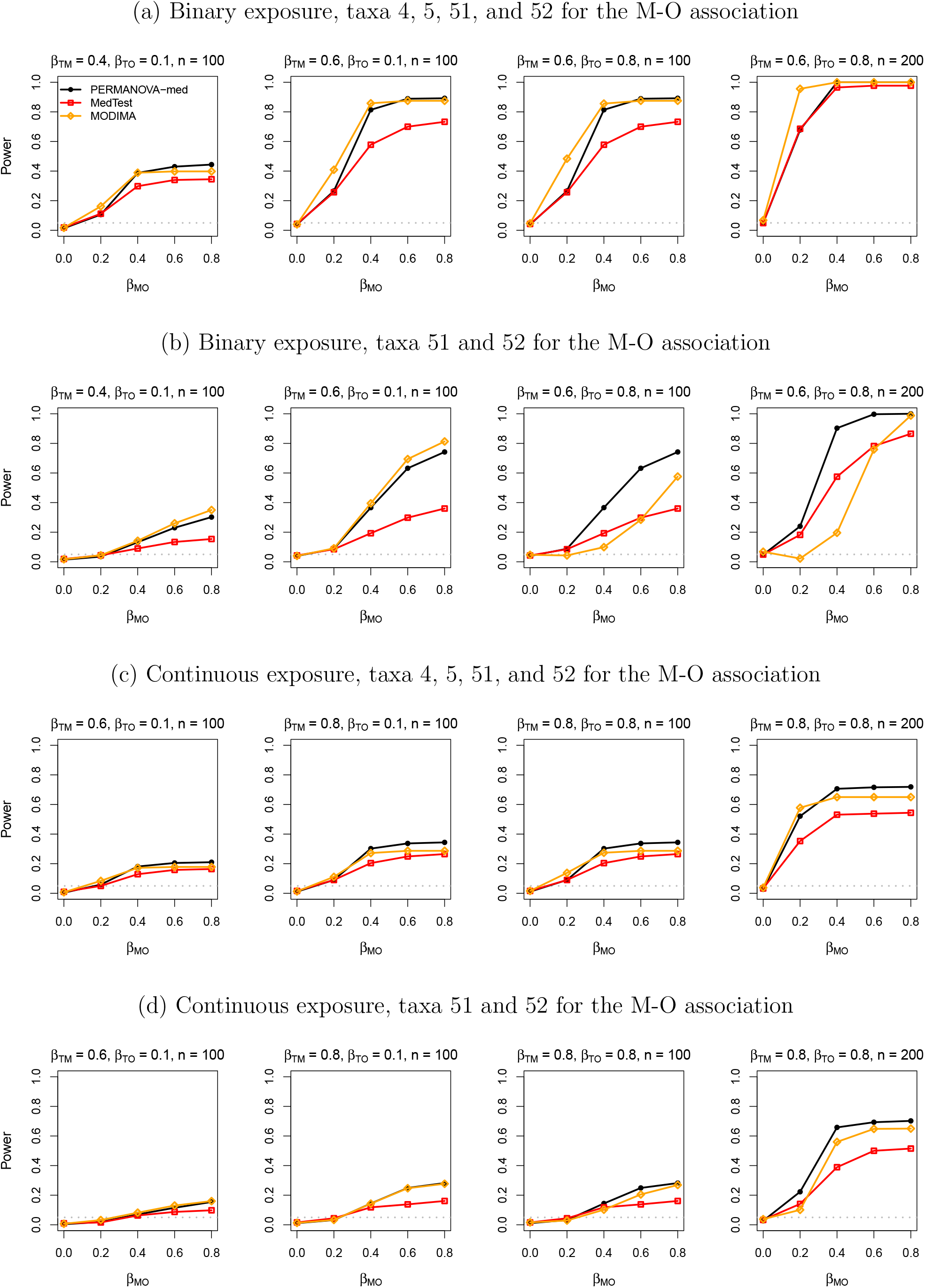
Simulation results in analysis of simulated data under M-mixed.

**Figure 5:**
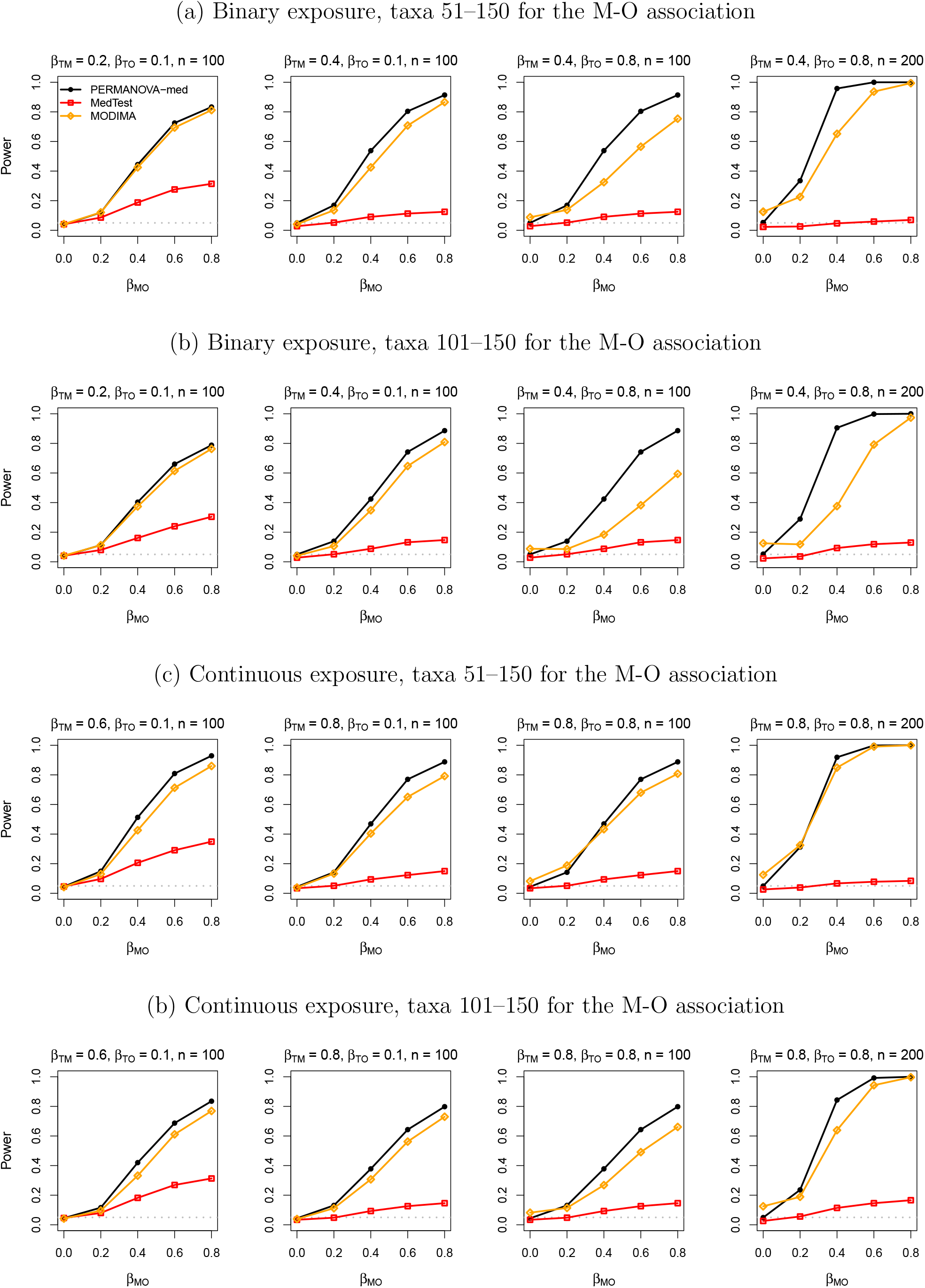
Simulation results in analysis of simulated data under M-rare.

Finally, when a confounder was added to the simulated data, MODIMA, without the capability to adjust for the confounding effect, produced very inflated type I error (Table 2). Note that, PERMANOVA-med and MedTest always controlled the type I error below the nominal level, with (Table 2) or without (Table 1) the confounder.

**Table 2:** Type I error (at the level 0.05) in analysis of simulated data with a binary exposure and a binary confounder

| Scenario | $\beta_{\text{TM}}$ | PERMANOVA-med | MedTest | MODIMA |
| --- | --- | --- | --- | --- |
| M-common | 0.2 | 0.008 | 0.014 | 0.242 |
|  | 0.4 | 0.035 | 0.040 | 0.385 |
| M-mixed | 0.4 | 0.006 | 0.015 | 0.056 |
|  | 0.6 | 0.020 | 0.029 | 0.103 |
| M-rare | 0.2 | 0.026 | 0.036 | 0.279 |
|  | 0.4 | 0.046 | 0.035 | 0.238 |

Note: $\beta_{\text{TO}} = 0.1$ , $\beta_{\text{MO}} = 0$ , and $n = 100$ . MODIMA does not allow adjustment of confounders.

### Real data on melanoma immunotherapy response

The real data [31] we used were generated from a cohort of 167 melanoma patients, who received immune checkpoint blockade (ICB) treatment and were classified as 106 responders and 61 non-responders. Their progression-free survival times (in days) were observed for 61 patients, censored for 49 patients, and missing for 57 patients. Their gut microbiome were profiled via shotgun metagenomic sequencing to generate a taxa count table including 225 taxa (lowest taxon known for a feature, up to species). These patients were further asked to complete a lifestyle survey, which included assessment of dietary fiber intake and use of probiotic supplements within the past month; 110 provided data for probiotic use, 94 provided data for dietary fiber intake, and 89 provided data for both.

Spencer et al. [31] found in this dataset that higher dietary fiber intake was associated with significantly improved progression-free survival, with the most pronounced benefit observed in patients with sufficient dietary fiber intake and no probiotic use. They also found marginal significance for the association of dietary fiber intake and response to ICB. In addition, the influence of the gut microbiome on immunotherapy response has been demonstrated in numerous human cohorts as well as in preclinical models [2, 32], and the human gut microbiome is itself shaped by diet [3] and medication use [33]. Given this interplay between diet and medication use, gut microbiome, and immunotherapy response, a natural question that arose was then whether some effect of dietary fiber intake and probiotic use on immunotherapy response in this dataset was mediated through the gut microbiome.

We performed a variety of mediation analyses using this dataset. For the outcome, we considered both the progression-free survival and the response to ICB, the former of which is a possibly censored survival time variable and the latter is a binary variable. For the exposure, we considered the dietary fiber intake (sufficient or insufficient), the probiotic use (no/yes), and the four-level categorical variable defined by both dietary fiber intake and probiotics use. Following [31], we additionally compared patients with sufficient dietary fiber intake and no probiotic use to all other three groups. We selected body mass index (BMI), prior treatment, lactate dehydrogenase level (LDH), and stage as potential confounders based on our analysis of bivariate associations, and we performed each mediation analyse with and without adjustment of these confounders. In all 16 mediation analyses, we applied PERMANOVA-med, MedTest, and MODIMA whenever they were applicable. For each method, we constructed tests based on the Bray-Curtis and Jaccard distance measures separately, as well as the omnibus test of both distance measures (except for MODIMA).

All results of *p*-values were summarized in Table 3. None of the *p*-values were significant at the 0.05 level, possibly due to the small sample sizes. Nevertheless, Table 3 demonstrated the wide applicability of PERMANOVA-med and the limited capabilities of MedTest and MODIMA. Specifically, neither MedTest nor MODIMA can handle censored survival times (the progression-free survival); MODIMA cannot adjust confounders (BMI et al.) nor provide an omnibus test (that combines Bray-Curtis and Jaccard); MedTest cannot handle multivariate exposures (the four-level categorical variable).

**Table 3:** *P*-values from 16 mediation analyses of the data on melanoma immunotherapy response

| Outcome | Exposure | n | PERMANOVA-med |  |  | MedTest |  |  | MODIMA |  |  |
| --- | --- | --- | --- | --- | --- | --- | --- | --- | --- | --- | --- |
|  |  |  | BC | J | Omni | BC | J | Omni | BC | J | Omni |
| No adjustment of covariates |  |  |  |  |  |  |  |  |  |  |  |
| Progression-free survival | Fiber intake | 89 | 0.808 | 0.965 | 0.958 | - | - | - | - | - | - |
|  | Probiotics | 110 | 0.913 | 0.716 | 0.899 | - | - | - | - | - | - |
|  | Fiber + probiotics (4 levels) | 89 | 0.777 | 0.975 | 0.953 | - | - | - | - | - | - |
|  | Sufficient fiber + no probiotics | 89 | 0.717 | 0.965 | 0.910 | - | - | - | - | - | - |
| Response to ICB | Fiber intake | 94 | 0.727 | 0.955 | 0.903 | 0.624 | 0.636 | 0.837 | 0.384 | 0.935 | - |
|  | Probiotics | 110 | 0.888 | 0.589 | 0.794 | 0.978 | 0.698 | 0.898 | 0.915 | 0.381 | - |
|  | Fiber + probiotics (4 levels) | 89 | 0.620 | 0.980 | 0.827 | - | - | - | 0.430 | 0.947 | - |
|  | Sufficient fiber + no probiotics | 89 | 0.490 | 0.955 | 0.697 | 0.276 | 0.626 | 0.455 | 0.441 | 0.947 | - |
| Adjusting for BMI, prior treatment, LDH, stage |  |  |  |  |  |  |  |  |  |  |  |
| Progression-free survival | Fiber intake | 89 | 0.786 | 0.990 | 0.936 | - | - | - | - | - | - |
|  | Probiotics | 110 | 0.983 | 0.788 | 0.947 | - | - | - | - | - | - |
|  | Fiber + probiotics (4 levels) | 89 | 0.770 | 0.995 | 0.935 | - | - | - | - | - | - |
|  | Sufficient fiber + no probiotics | 89 | 0.725 | 0.980 | 0.903 | - | - | - | - | - | - |
| Response to ICB | Fiber intake | 94 | 0.870 | 0.920 | 0.975 | 0.832 | 0.935 | 0.966 | - | - | - |
|  | Probiotics | 110 | 0.973 | 0.433 | 0.630 | 0.911 | 0.539 | 0.773 | - | - | - |
|  | Fiber + probiotics (4 levels) | 89 | 0.760 | 0.975 | 0.928 | - | - | - | - | - | - |
|  | Sufficient fiber + no probiotics | 89 | 0.644 | 0.925 | 0.850 | 0.453 | 0.973 | 0.682 | - | - | - |

Note: BC: Bray-Curtis; J: Jaccard; Omni: the omnibus test that combines the results from analyzing the Bray-Curtis and Jaccard distances; $n$ : sample size; -: not applicable.

## Discussion

We presented PERMANOVA-med, an extension of PERMANOVA to mediation analysis of microbiome data. Through extensive simulation studies, we observed that PERMANOVA-med did not uniformly outperform MedTest. However, the scenarios in which PERMANOVA-med did outperform seemed more realistic and more general, e.g., scenarios with a mixture of abundant and less abundant mediating taxa, relatively rare mediating taxa, or different sets of taxa associated with the exposure and the outcome. Even in the single scenario that PERMANOVA-med lost power to MedTest (Figure 3(a)), the power loss was relatively small. The power comparison between PERMANOVA-med and MODIMA was more difficult, as MODIMA often lost control of the type I error. Nevertheless, there were many more scenarios in which PERMANOVA-med had higher power than MODIMA than scenarios when it was the opposite.

The main advantage of PERMANOVA-med over MedTest and MODIMA is its wide applicability to a variety of mediation analyses of microbiome data, which was achieved by using our existing function permanovaFL. Through analysis of the simulated data and the real data, we have illustrated most features in Figure 2 that are supported by permanovaFL, such as multivariate exposures, survival outcomes, and omnibus tests of multiple distance measures. Although we did not cover clustered or matched-set data in this article, these types of data are emerging rapidly in recent years and may also call for mediation analysis. PERMANOVA-med is well positioned to accommodate such data in its current form.

## References

1. Bai J, Hu Y, Bruner D. Composition of gut microbiota and its association with body mass index and lifestyle factors in a cohort of 7–18 years old children from the American Gut Project. Pediatric Obesity. 2019;14(4):e12480.

2. Routy B, Le Chatelier E, Derosa L, Duong CP, Alou MT, Daillére R, et al. Gut microbiome influences efficacy of PD-1–based immunotherapy against epithelial tumors. Science. 2018;359(6371):91–97.

3. McDonald D, Hyde E, Debelius JW, Morton JT, Gonzalez A, Ackermann G, et al. American gut: an open platform for citizen science microbiome research. Msystems. 2018;3(3):e00031–18.

4. Hu YJ, Satten GA. Testing hypotheses about the microbiome using the linear decomposition model (LDM). Bioinformatics. 2020;p. bbtaa260, https://doi.org/10.1093/bioinformatics/btaa260.

5. Zhu Z, Satten GA, Mitchell C, Hu YJ. Constraining PERMANOVA and LDM to within-set comparisons by projection improves the efficiency of analyses of matched sets of microbiome data. Microbiome. 2021;9(1):1–19.

6. Legendre P, Anderson MJ. Distance-based redundancy analysis: testing multispecies responses in multifactorial ecological experiments. Ecological monographs. 1999;69(1):1– 24.

7. McArdle BH, Anderson MJ. Fitting multivariate models to community data: a comment on distance-based redundancy analysis. Ecology. 2001;82(1):290–297.

8. Zhao N, Chen J, Carroll IM, Ringel-Kulka T, Epstein MP, Zhou H, et al. Testing in microbiome-profiling studies with MiRKAT, the microbiome regression-based kernel association test. The American Journal of Human Genetics. 2015;96(5):797–807. PMCID: PMC4570290.

9. Alekseyenko AV. Multivariate Welch t-test on distances. Bioinformatics. 2016;32(23):3552–3558.

10. Zhang Y, Han SW, Cox LM, Li H. A multivariate distance-based analytic framework for microbial interdependence association test in longitudinal study. Genetic epidemiology. 2017;41(8):769–778.

11. Jaccard P. The distribution of the flora in the alpine zone. 1. New phytologist. 1912;11(2):37–50.

12. Bray JR, Curtis JT. An ordination of the upland forest communities of southern Wisconsin. Ecological monographs. 1957;27(4):326–349.

13. Lozupone C, Knight R. UniFrac: a new phylogenetic method for comparing microbial communities. Applied and environmental microbiology. 2005;71(12):8228–8235. PMCID: PMC1317376.

14. Chen J, Bittinger K, Charlson ES, Hoffmann C, Lewis J, Wu GD, et al. Associating microbiome composition with environmental covariates using generalized UniFrac distances. Bioinformatics. 2012;28(16):2106–2113.

15. Zhang J, Wei Z, Chen J. A distance-based approach for testing the mediation effect of the human microbiome. Bioinformatics. 2018;34(11):1875–1883.

16. Hamidi B, Wallace K, Alekseyenko AV. MODIMA, a Method for Multivariate Omnibus Distance Mediation Analysis, Allows for Integration of Multivariate Exposure-Mediator-Response Relationships. Genes. 2019;10(7):524.

17. Székely GJ, Rizzo ML, Bakirov NK. Measuring and testing dependence by correlation of distances. The annals of statistics. 2007;35(6):2769–2794.

18. Székely GJ, Rizzo ML. Brownian distance covariance. The annals of applied statistics. 2009;3(4):1236–1265.

19. Székely GJ, Rizzo ML. Partial distance correlation with methods for dissimilarities. The Annals of Statistics. 2014;42(6):2382–2412.

20. Hu Y, Satten GA, Hu YJ. Testing microbiome associations with censored survival outcomes at both the community and individual taxon levels. bioRxiv. 2022;.

21. Tang ZZ, Chen G, Alekseyenko AV. PERMANOVA-S: association test for microbial community composition that accommodates confounders and multiple distances. Bioinformatics. 2016;32(17):2618–2625.

22. Hu YJ, Satten GA. A rarefaction-without-resampling extension of PERMANOVA for testing presence-absence associations in the microbiome. bioRxiv. 2021;p. https://doi.org/10.1101/2021.04.06.438671.

23. Baron RM, Kenny DA. The moderator–mediator variable distinction in social psychological research: Conceptual, strategic, and statistical considerations. Journal of personality and social psychology. 1986;51(6):1173.

24. VanderWeele TJ, Vansteelandt S. Conceptual issues concerning mediation, interventions and composition. Statistics and its Interface. 2009;2(4):457–468.

25. O’Reilly PF, Hoggart CJ, Pomyen Y, Calboli FC, Elliott P, Jarvelin MR, et al. MultiPhen: joint model of multiple phenotypes can increase discovery in GWAS. PloS One. 2012;7(5).

26. Wu B, Pankow JS. Statistical methods for association tests of multiple continuous traits in genome-wide association studies. Annals of Human Genetics. 2015;79(4):282–293.

27. Majumdar A, Witte JS, Ghosh S. Semiparametric allelic tests for mapping multiple phenotypes: Binomial regression and Mahalanobis distance. Genetic Epidemiology. 2015;39(8):635–650.

28. Gower JC. Some distance properties of latent root and vector methods used in multivariate analysis. Biometrika. 1966;53(3-4):325–338.

29. Freedman D, Lane D. A nonstochastic interpretation of reported significance levels. Journal of Business & Economic Statistics. 1983;1(4):292–298.

30. Charlson ES, Chen J, Custers-Allen R, Bittinger K, Li H, Sinha R, et al. Disordered microbial communities in the upper respiratory tract of cigarette smokers. PloS one. 2010;5(12):e15216. PMCID: PMC3004851.

31. Spencer CN, McQuade JL, Gopalakrishnan V, McCulloch JA, Vetizou M, Cogdill AP, et al. Dietary fiber and probiotics influence the gut microbiome and melanoma immunotherapy response. Science. 2021;374(6575):1632–1640.

32. Matson V, Fessler J, Bao R, Chongsuwat T, Zha Y, Alegre ML, et al. The commensal microbiome is associated with anti–PD-1 efficacy in metastatic melanoma patients. Science. 2018;359(6371):104–108.

33. Maier L, Pruteanu M, Kuhn M, Zeller G, Telzerow A, Anderson EE, et al. Extensive impact of non-antibiotic drugs on human gut bacteria. Nature. 2018;555(7698):623– 628.

